# Hyper-Hierarchical Brain States Are Associated with Disorders of Consciousness

**DOI:** 10.64898/2026.07.30.741772

**Authors:** K Olaciregui-Dague, I Acero-Pousa, T Berjaga-Buisan, M Kaller, JD Sitt, G Deco, ML Kringelbach

## Abstract

**Background:** Consciousness is increasingly understood as an emergent property of large-scale brain dynamics that depend upon flexible interactions among distributed cortical and subcortical systems. Although disorders of consciousness (DOC) have traditionally been associated with impaired integration and reduced network complexity, the role of hierarchical brain organization in supporting conscious awareness remains poorly understood. Here, we investigated how hierarchical organization relates to behavioral responsiveness in DOC by combining trophic-level analysis, trophic coherence, and whole-brain dynamical metrics.

**Methods:** Resting-state functional MRI data were analyzed from healthy controls (CNT), minimally conscious state (MCS) patients, and unresponsive wakefulness syndrome (UWS) patients drawn from a previously published DOC cohort. Static global and regional measures of functional hierarchy were computed from directed effective-connectivity networks. Dynamic trophic states were identified using time-resolved phase-coupling analyses and clustering of recurrent coordination patterns. State occupancy, dwell time, metastability, synchrony, and behavioral associations with Coma Recovery Scale–Revised (CRS-R) scores were evaluated.

**Results:** Regional trophic levels were positively associated with behavioral responsiveness, with higher frontal and thalamic trophic levels and lower insular trophic levels predicting higher Coma Recovery Scale–Revised (CRS-R) scores. Dynamic trophic-state analysis identified a pathological hyper-hierarchical state, defined by elevated frontal, thalamic, and insular trophic levels, that exhibited progressively greater occupancy and longer dwell times from healthy controls to minimally conscious state and unresponsive wakefulness syndrome patients. In contrast, occupancy and dwell time of this state distinguished diagnostic groups but were not significantly associated with behavioral responsiveness. Independent analyses demonstrated significant reductions in metastability and global synchrony across disorders of consciousness. Anatomical mapping localized elevated trophic levels within the pathological state predominantly to fronto-thalamo-limbic systems.

**Conclusions:** Disorders of consciousness are characterized not simply by loss of hierarchical organization but by prolonged stabilization within recurrent hyper-hierarchical brain states. Conscious awareness appears to depend not only on hierarchical organization itself but also on the capacity to flexibly transition between distinct brain states. Severe disorders of consciousness are associated with persistent occupation of pathological hyper-hierarchical states, potentially restricting the dynamical repertoire available for conscious processing.

## Introduction

Consciousness emerges from the coordinated activity of distributed neural systems operating across multiple spatial and temporal scales. Despite substantial advances in cognitive neuroscience, understanding how conscious awareness arises from the large-scale hierarchical organization of the brain remains one of the central challenges of modern neuroscience. Disorders of consciousness (DOC), including minimally conscious state (MCS) and unresponsive wakefulness syndrome (UWS), provide a unique opportunity to investigate this problem because they encompass varying degrees of preserved wakefulness and awareness following severe brain injury^1–4^.

Over the last two decades, converging evidence from neuroimaging, electrophysiology, and computational modelling has demonstrated that conscious awareness depends upon interactions among widely distributed cortical and subcortical systems rather than activity within any single anatomical region^5–9^. Global Neuronal Workspace Theory proposes that consciousness emerges when information becomes globally available through recurrent frontoparietal broadcasting mechanisms^10,11^. Integrated Information Theory similarly emphasizes large-scale integration while highlighting the importance of informational differentiation^12^. More recently, dynamical systems approaches have argued that consciousness depends upon the capacity of the brain to flexibly transition among multiple metastable brain states rather than remaining confined to a single stable configuration^13–16^.

Consistent with these perspectives, disorders of consciousness are associated with widespread disruptions of functional connectivity, thalamocortical communication, and large-scale integration^17–21^. Functional MRI studies have repeatedly demonstrated abnormalities within the default mode network, salience network, frontoparietal control systems, and thalamocortical circuits in DOC patients^22–25^. However, increasing evidence suggests that static connectivity alone provides an incomplete description of conscious brain function. Resting-state activity exhibits substantial temporal variability, continuously transitioning among recurring brain states that differ in their patterns of integration and segregation^26–29^. Whether hierarchical organization itself dynamically reorganizes across these recurring brain states—and whether such reorganization is altered in disorders of consciousness—remains unknown.

The importance of these dynamic processes has become increasingly apparent in studies of consciousness. Dynamic functional connectivity analyses have shown that healthy conscious brains explore a rich repertoire of transient brain states, whereas reduced consciousness is associated with diminished state diversity and altered transition dynamics^30–33^. In patients with DOC, conscious awareness has been linked to the preservation of flexible network reconfiguration and the capacity to explore multiple dynamical regimes^34^. Together, these findings suggest that consciousness depends not merely on the presence of specific network structures but on the capacity to flexibly transition among distinct brain states.

One influential framework for characterizing such flexibility arises from whole-brain computational modelling. Using coupled oscillator models and empirical neuroimaging data, Deco, Kringelbach, Cabral, and colleagues have proposed that healthy brain function operates within a metastable regime characterized by continual fluctuations between integrated and segregated brain states^14,15,45,62^. Within this framework, global synchrony reflects the average degree of coordination across the brain, whereas metastability quantifies the temporal variability of that coordination. Conscious states are typically associated with elevated metastability, reflecting an optimal balance between stability and flexibility, whereas unconscious states exhibit reduced dynamical complexity and restricted exploration of state space^18,32,64,66^.

Although these approaches have yielded important insights into the dynamics of consciousness, comparatively little attention has been paid to the role of directed network organization. Neural systems exhibit pronounced hierarchical organization spanning sensory, associative, limbic, and integrative systems^35–37^. Hierarchical organization has long been recognized as a fundamental principle of brain architecture, influencing information flow, predictive processing, and large-scale coordination^38–40^. However, most studies of hierarchy in neuroscience have focused on anatomical connectivity or laminar organization rather than directed interactions inferred from whole-brain functional dynamics.

Trophic-level analysis provides a graph-theoretic framework for quantifying hierarchical organization within directed networks. Originally developed for ecological systems, trophic levels estimate the relative position of nodes within directed interaction networks, whereas trophic coherence quantifies the consistency of directed interactions with this trophic ordering^41,42^. Recent applications have demonstrated the utility of trophic approaches for characterizing complex directed systems, yet their relevance to consciousness remains largely unexplored. Nevertheless, recent work using an alternative hierarchy framework has likewise implicated disrupted hierarchical organization in disorders of consciousness, suggesting that hierarchical organization may represent a general feature of conscious brain dynamics.^49^

We hypothesized that hierarchical organization may exhibit fundamentally different relationships with consciousness across static and dynamic timescales. Higher trophic levels within fronto-thalamic systems were expected to support residual conscious processing, whereas excessive concentration of hierarchical organization within recurrent dynamic brain states was hypothesized to reflect pathological rigidity and reduced access to a rich dynamical repertoire.

To test this hypothesis, we combined trophic network analysis, dynamic trophic state analysis, and whole-brain dynamical metrics in a cohort of healthy controls, minimally conscious state patients, and unresponsive wakefulness syndrome patients. First, we quantified regional trophic levels and global trophic coherence within directed effective connectivity networks. Second, we identified recurrent dynamic trophic states and characterized their occupancy, dwell time, and behavioral relevance. Third, we quantified metastability and global synchrony using established whole-brain dynamical frameworks. Finally, we mapped the anatomical distribution of state-specific trophic levels and evaluated their relationship to behavioral responsiveness.

We demonstrate a striking dissociation between static and dynamic hierarchical organization. Whereas higher frontal and thalamic trophic levels within static effective-connectivity networks predict better behavioral responsiveness, recurrent dynamic trophic states associated with disorders of consciousness are characterized by elevated regional trophic levels together with increased occupancy and prolonged dwell times. These pathological dynamics are accompanied by reduced metastability and altered global synchrony. Together, these findings suggest that consciousness depends not simply on hierarchical organization itself, but on the capacity to flexibly transition among distinct hierarchical brain states.

## Methods

### Participants

Participants were drawn from the publicly available disorders of consciousness (DOC) dataset originally described by Sitt and colleagues^31^. This dataset has served as the basis for multiple studies investigating the large-scale neural dynamics of consciousness.^2–4^ The final cohort comprised 13 healthy controls (7 females; mean age 42.5 ± 13.6 years), 11 patients in the minimally conscious state (5 females; mean age 47.3 ± 20.8 years), and 10 patients with unresponsive wakefulness syndrome (4 females; mean age 39.3 ± 16.3 years).

Clinical diagnosis was established using repeated behavioral assessments and standardized diagnostic criteria for disorders of consciousness^2,4^. Behavioral responsiveness was quantified using the Coma Recovery Scale–Revised (CRS-R), which remains the most widely validated instrument for assessing residual consciousness following severe brain injury^44^.

### MRI Acquisition and Regional Time Series

Resting-state functional MRI data were acquired and preprocessed according to the procedures described in the original dataset publication^31^. Regional mean blood oxygen level-dependent (BOLD) time series were extracted from the 90 cortical and subcortical regions of the Automated Anatomical Labeling (AAL90) atlas and used in all subsequent analyses^43^.

### Structural Connectivity

A structural connectivity (SC) matrix was used as the anatomical scaffold for whole-brain dynamical modelling and effective connectivity estimation. The matrix comprised weighted inter-regional anatomical connections among the 90 regions of the AAL90 atlas and was normalized to unit maximum weight prior to model fitting. Rather than being analyzed directly, the SC matrix provided the fixed anatomical architecture constraining interactions among brain regions during model optimization. This approach ensured that estimated effective connectivity remained biologically plausible while allowing participant-specific directed interactions to emerge from the empirical resting-state functional data.

### Whole-brain dynamical modelling

Participant-specific effective connectivity was estimated using a generative whole-brain model based on coupled Hopf oscillators. In this framework, each of the 90 AAL brain regions was represented by a nonlinear oscillator operating near the Hopf bifurcation, enabling realistic simulation of resting-state brain dynamics while maintaining computational efficiency.

Oscillators were coupled according to the empirical structural connectivity matrix, which provided the fixed anatomical scaffold for large-scale interactions. Empirical resting-state functional connectivity and time-lagged covariance matrices derived from the regional BOLD time series were used as optimization targets. Model parameters were iteratively adjusted until the simulated dynamics reproduced the empirical functional organization while remaining constrained by the underlying structural architecture.

Optimization yielded participant-specific directed effective connectivity (EC) matrices describing the strength and direction of interactions between all brain regions. These EC matrices formed the basis for all subsequent trophic hierarchy, trophic coherence, and dynamic state analyses.

### Effective Connectivity Estimation

The optimized effective connectivity matrices were represented as weighted directed graphs,

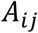

where *A_ij_* denotes the strength of the directed influence from brain region *i* to brain region *j*.

### Trophic levels

Hierarchical organization was quantified using trophic levels, a graph-theoretic framework first developed in ecological systems^41,42^ and subsequently adapted to characterize directed organization in complex systems.

Trophic levels were computed according to:

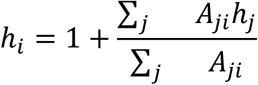

where ℎ*_i_*denotes the trophic level of node *i*, and *A_ij_* represents the weight of the directed connection from node *j* to node *i*.

Trophic levels quantify the relative hierarchical position of each node within a directed network, providing a principled measure of hierarchical organization that is independent of simple degree-based metrics^41,42,49–50^. Higher trophic levels indicate progressively more downstream positions within the directed network hierarchy. Trophic levels were estimated using graph Laplacian methods and the Moore–Penrose pseudoinverse, following established trophic-network approaches^41,42^.

### Trophic Coherence

Global hierarchical organization was quantified using trophic coherence, which measures the consistency of directed interactions with the inferred hierarchy^41, 42^.

For each directed edge:

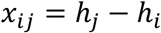

where *x_ij_* denotes the trophic difference between connected nodes. Trophic incoherence (*q*) was computed from the variance of trophic differences across all directed edges, and trophic coherence was defined as

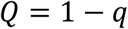

where larger values indicate stronger hierarchical organization.

### Dynamic Brain-State Identification

Time-varying brain dynamics were characterized using a dynamic phase-coupling framework based on Leading Eigenvector Dynamics Analysis (LEiDA)^27^. This approach identifies recurrent large-scale patterns of phase synchronization from resting-state BOLD signals, enabling characterization of transient brain states over time.

### Phase Extraction

Regional BOLD time series were band-pass filtered to the canonical resting-state frequency range (0.008–0.08 Hz) and transformed into instantaneous phase representations using the Hilbert transform^45,68^.

The analytic signal was defined as

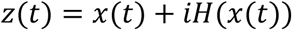

and instantaneous phase was obtained as

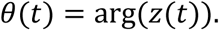

This procedure follows the phase-based framework introduced by Cabral *et al.*^45^ and subsequently adopted in studies of whole-brain dynamics, metastability, and disorders of consciousness^15,30,32,65^.

### Dynamic Phase Coupling

At each time point, instantaneous phase-locking between all pairs of brain regions was quantified as

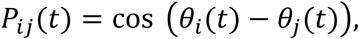

yielding time-resolved phase-locking matrices describing the instantaneous large-scale synchronization of regional brain activity^25,29,32^.

The leading eigenvector of each phase-locking matrix was subsequently extracted to characterize the dominant large-scale pattern of phase synchronization at each time point. These leading eigenvectors formed the input for dynamic state identification using Leading Eigenvector Dynamics Analysis (LEiDA).

### Dynamic State Identification

Recurring brain states were identified using k-means clustering of the leading eigenvectors obtained from the instantaneous phase-locking matrices. The optimal number of clusters was evaluated using elbow analysis of the within-cluster sum of squares (Supplementary Fig. 1). Because the elbow curve exhibited diminishing reductions beyond approximately *k* = 5–6 rather than a single well-defined optimum, clustering solutions ranging from *k* = 5 to *k* = 9 were examined. A seven-state solution was selected for the primary analyses because it provided a parsimonious and biologically interpretable decomposition of the hierarchy dynamics while maintaining clear separation between the healthy/integrative and pathological/UWS-enriched trophic states. For each participant, every time point was assigned to one of the seven recurrent states, enabling quantification of state-specific fractional occupancy and mean dwell time. The robustness of the principal findings across alternative clustering resolutions was evaluated in complementary analyses (see Results, Robustness across clustering resolutions).

### State-specific Effective Connectivity

Effective connectivity matrices were estimated for each participant throughout the resting-state scan. Following LEiDA clustering, effective connectivity matrices corresponding to time points assigned to the same dynamic state were averaged to generate state-specific effective connectivity networks. Regional trophic levels and global trophic coherence were subsequently computed from these state-specific networks.

### Dynamic State Metrics

For each participant and dynamic state, fractional occupancy and mean dwell time were computed. Fractional occupancy was defined as the proportion of time points assigned to a given state, whereas mean dwell time quantified the average duration of consecutive visits to that state.

State-specific regional trophic levels (frontal, thalamic, and insular) and global trophic coherence were subsequently computed from the corresponding state-specific effective connectivity networks. These measures were compared across diagnostic groups and correlated with behavioral responsiveness, as assessed by the Coma Recovery Scale–Revised (CRS-R).

### Metastability and Synchrony

To independently characterize whole-brain dynamics, synchrony and metastability were quantified using the phase-based framework developed by Cabral, Deco, Kringelbach, and colleagues^25–31^.

#### Kuramoto Order Parameter

Global synchronization was quantified using the Kuramoto order parameter,

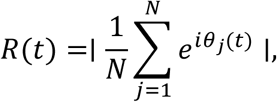

where *N* is the number of brain regions, *θ_i_*(*t*) is the instantaneous phase of region *j* at time *t*, *i* is the imaginary unit, and ∣⋅∣ denotes the magnitude of the mean phase vector. *R*(*t*) ranges from 0 (complete phase dispersion) to 1 (perfect phase synchronization), providing a measure of instantaneous whole-brain synchronization.

#### Synchrony

Mean synchrony was defined as the temporal average of the Kuramoto order parameter,

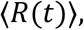

representing the average level of whole-brain synchronization throughout the resting-state scan^25–27^.

#### Metastability

Metastability was quantified as the temporal standard deviation of the Kuramoto order parameter,

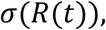

providing a measure of the temporal variability of whole-brain synchronization^27,29^.

### Anatomical Visualization

Regional trophic-level values were projected onto the AAL90 atlas and visualized using Nilearn^47^ to generate cortical surface renderings. State-specific trophic-level maps and contrast maps were generated to illustrate the anatomical distribution of hierarchical organization across dynamic brain states.

### Statistical Analysis

Group differences were assessed using one-way analysis of variance (ANOVA), followed by Tukey’s honestly significant difference post hoc tests where appropriate. Associations between trophic metrics and behavioral responsiveness were evaluated using Spearman rank correlations. The joint contribution of regional trophic levels to behavioral responsiveness was assessed using multiple linear regression. Effect sizes for ANOVA are reported as eta-squared (η^2^). Statistical significance was defined as a two-sided *P* < 0.05. All analyses were performed in MATLAB R2025b (MathWorks, Natick, MA, USA) unless otherwise specified. As a sensitivity analysis, the multiple regression relating regional trophic levels to CRS-R was repeated with participant age included as a covariate, and model stability was assessed using leave-one-out re-estimation (Supplementary Table 2, Supplementary Fig. 4).

### Hidden Markov Model Sensitivity Analysis

To evaluate whether the pathological/UWS-enriched dynamic trophic state constitutes a robust feature of DOC dynamics rather than an artefact of the LEiDA/k-means clustering pipeline, we performed a convergent-validity analysis using an independent dynamic-state framework. A Gaussian hidden Markov model (HMM) was fitted directly to the band-pass filtered, z-scored regional BOLD time series using the HMM-MAR toolbox^26,69^, without LEiDA’s leading-eigenvector dimensionality-reduction step. States were modelled as state-specific covariance structures with a zero-mean assumption (model order = 0; full covariance), consistent with prior applications of this framework to resting-state fMRI dynamics^26^. Models were fitted independently for *K* = 5–9 states, mirroring the robustness analysis performed for the LEiDA solution. For each *K*, the state with the highest mean fractional occupancy in healthy controls was designated the healthy-candidate state, and the state with the highest mean occupancy in patients with unresponsive wakefulness syndrome was designated the pathological-candidate state. For each subject, fractional occupancy, mean dwell time, and the empirical self-transition probability (the probability of remaining within a state at the subsequent timepoint, estimated from the Viterbi state path) were computed for these two states and compared across diagnostic groups (one-way ANOVA) and correlated with CRS-R (Spearman). Because occupancy of both states was strongly bimodal across subjects, with most subjects showing either negligible or substantial occupancy rather than intermediate values, we additionally tested whether the probability of ever occupying a given state (occupancy > 0) differed across diagnostic groups using a chi-squared test of independence.

## Results

### Regional trophic levels predict behavioral responsiveness

We first investigated whether trophic levels are altered in disorders of consciousness and whether these alterations relate to behavioral responsiveness (Fig. 2). Trophic coherence showed a progressive reduction from healthy controls (CNT; 0.008 ± 0.006) to minimally conscious state (MCS; 0.005 ± 0.003) and unresponsive wakefulness syndrome (UWS; 0.003 ± 0.002), although this trend did not reach statistical significance (one-way ANOVA, *F*(2,31) = 3.038, *P* = 0.062, η^2^ = 0.164).

**Figure 1.**
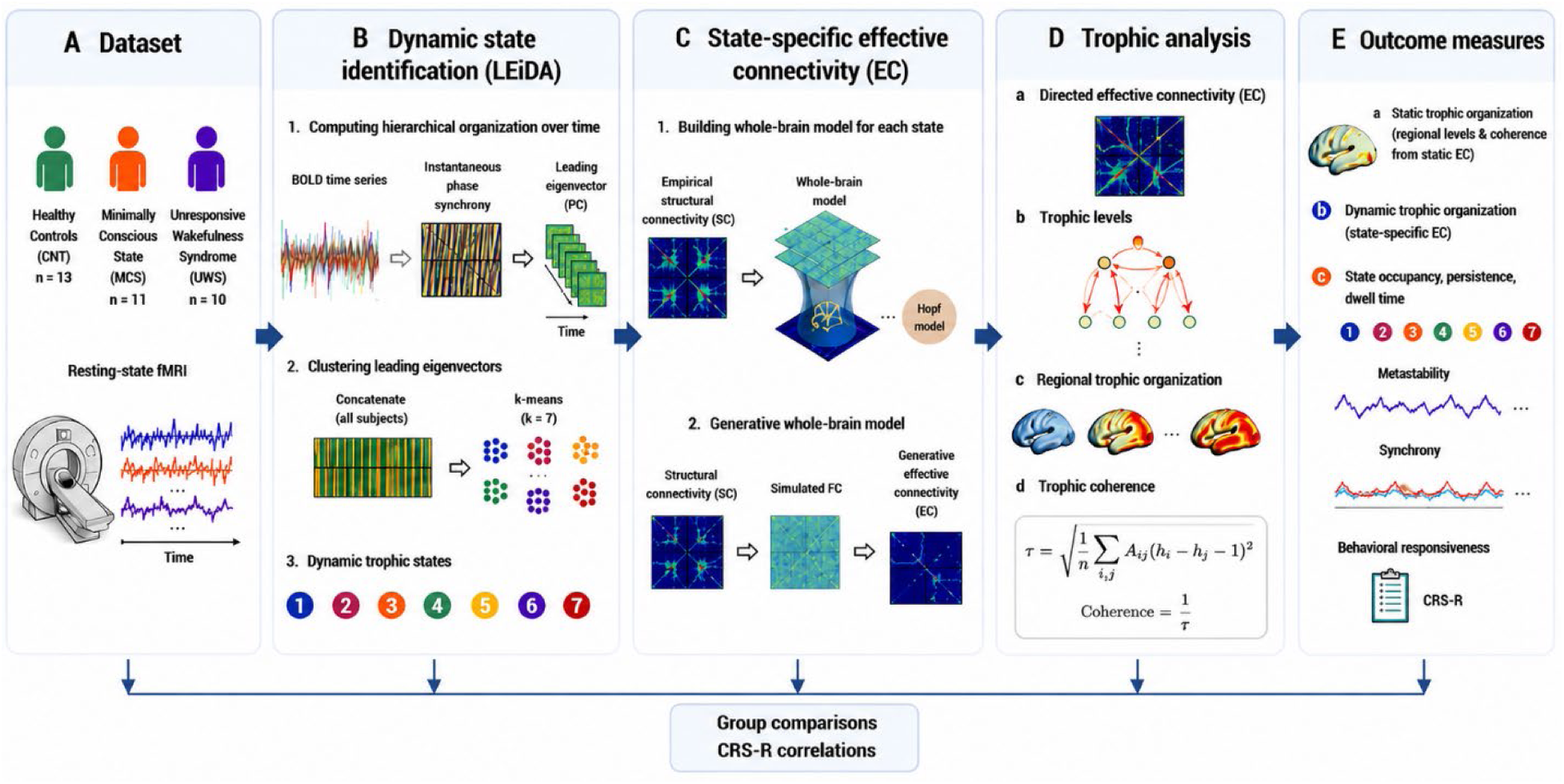
| Analytical workflow for identifying dynamic trophic organization in disorders of consciousness. Resting-state fMRI data were acquired from healthy controls (CNT, *n*=13), minimally conscious state (MCS, *n*=11), and unresponsive wakefulness syndrome (UWS, *n*=10) participants. Dynamic functional states were identified using Leading Eigenvector Dynamics Analysis (LEiDA). For each state, empirical structural and functional connectivity constrained whole-brain effective connectivity (EC) estimation using the Hopf whole-brain model. Directed EC matrices were analyzed using trophic network theory to derive regional trophic levels and global trophic coherence. Dynamic-state occupancy and dwell time, together with whole-brain dynamical measures (metastability and synchrony), were related to behavioral responsiveness (Coma Recovery Scale–Revised; CRS-R).

**Figure 2.**
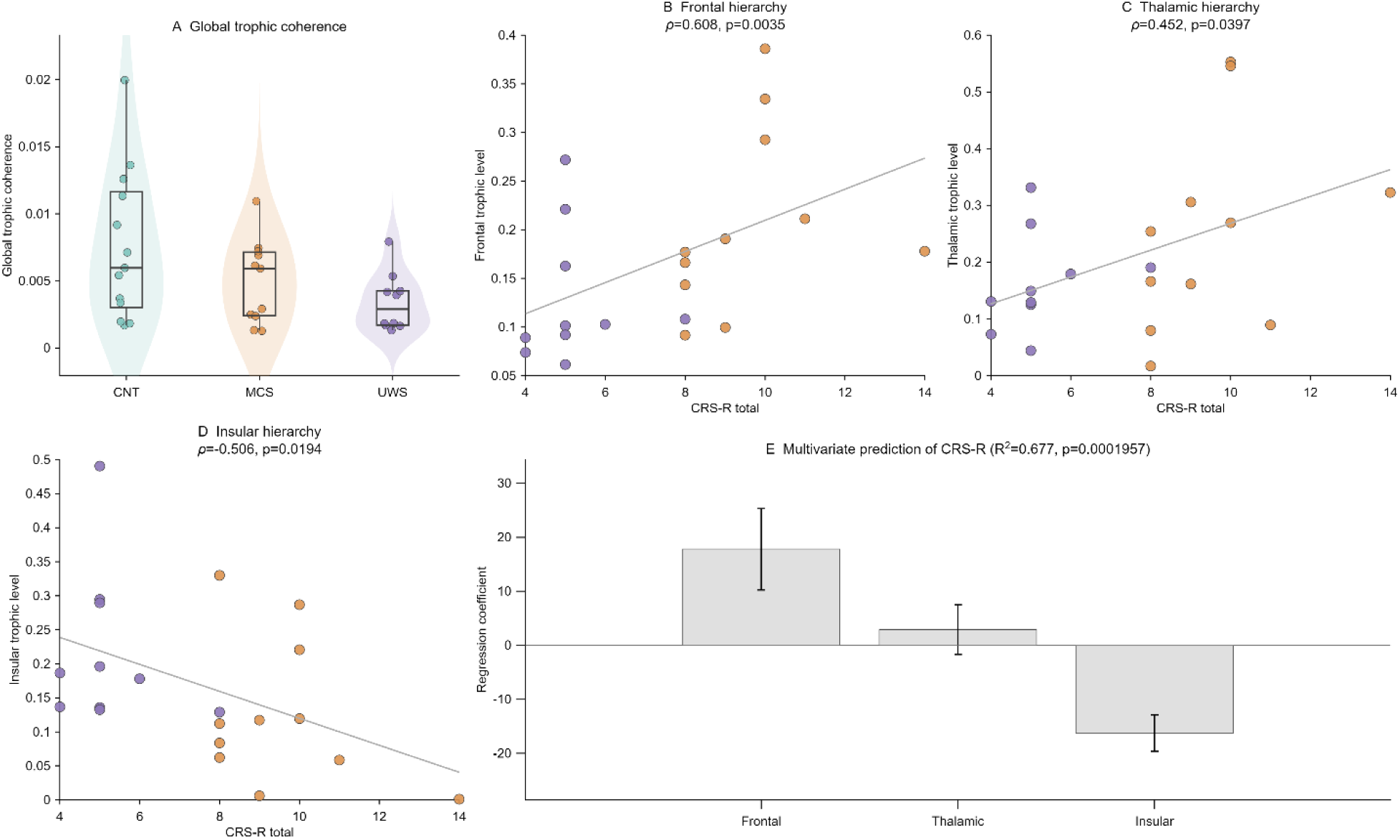
| Static trophic hierarchy predicts behavioral responsiveness. **a**, Global trophic coherence across groups. Although global trophic coherence tended to decrease from CNT to UWS, the omnibus group effect did not reach significance (one-way ANOVA, *F*(2,31)=3.04, *P*=0.062, η^2^=0.164). **b–d**, Regional trophic levels correlated with behavioral responsiveness. Frontal trophic level showed a strong positive association with CRS-R score (Spearman ρ=0.608, *P*=0.00348), thalamic trophic level showed a moderate positive association (ρ=0.452, *P*=0.0397), whereas insular trophic level was negatively associated with CRS-R (ρ=−0.506, *P*=0.0194). Regression lines indicate least-squares fits. **e**, Multivariate linear regression demonstrated that regional trophic levels jointly predicted behavioral responsiveness (R^2^=0.677, adjusted R^2^=0.620, overall model *P*=1.96×10–^4^). Bars denote unstandardized regression coefficients ± s.e.m.

Trophic levels revealed anatomically selective alterations concentrated within frontal association cortex rather than widespread disruption of cortical hierarchical organization. Four frontal regions exhibited uncorrected group differences (*P* < 0.05), including the left superior medial frontal cortex (*F*(2,31) = 3.644, *P* = 0.038, η^2^ = 0.190), right orbitomedial superior frontal cortex (*F*(2,31) = 4.407, *P* = 0.021, η^2^ = 0.221), right superior medial frontal cortex (*F*(2,31) = 4.965, *P* = 0.013, η^2^ = 0.243), and right superior frontal cortex (*F*(2,31) = 4.162, *P* = 0.025, η^2^ = 0.212). The largest group difference was observed in the right superior medial frontal cortex (CNT versus UWS, Cohen’s *d* = 1.257).

To determine whether regional trophic levels were associated with behavioral responsiveness, trophic levels were correlated with Coma Recovery Scale–Revised (CRS-R) scores across patients. The strongest positive associations were observed in the right superior medial frontal cortex (Spearman ρ = 0.661, *P* = 0.001), left superior medial frontal cortex (ρ = 0.610, *P* = 0.003), right supplementary motor area (ρ = 0.509, *P* = 0.018), and left orbitomedial superior frontal cortex (ρ = 0.489, *P* = 0.024), whereas the strongest negative association was identified in the left insula (ρ = −0.506, *P* = 0.019).

To evaluate these relationships at the systems level, trophic levels were averaged across frontal, thalamic, and insular regions. Frontal trophic level correlated positively with behavioral responsiveness (ρ = 0.608, *P* = 0.003), as did thalamic trophic level (ρ = 0.452, *P* = 0.040), whereas insular trophic level was negatively associated (ρ = −0.506, *P* = 0.019).

A multiple linear regression incorporating these three regional measures explained 67.7% of the variance in behavioral responsiveness (*R*^2^ = 0.677; adjusted *R*^2^ = 0.620; *F*(3,17) = 11.867; model *P* = 1.96 × 10–^4^). Frontal trophic level (β = 17.821, *P* = 0.030) and insular trophic level (β = −16.284, *P* = 1.53 × 10–^4^) remained independent predictors of CRS-R, whereas thalamic trophic level was not independently associated (β = 2.910, *P* = 0.536).

Because patient age varied considerably across the cohort, we repeated this analysis with age included as a covariate (Supplementary Table 2, Supplementary Fig. 4). Age did not improve model fit (ΔR^2^ = 0.003; F-change(1,16) = 0.157, *P* = 0.697) and was not itself associated with CRS-R (β = 0.010, *P* = 0.697). Frontal (β = 17.897, *P* = 0.034) and insular (β = −15.865, *P* = 4.50 × 10–^4^) trophic levels remained independent predictors of behavioral responsiveness after age adjustment, indicating that these associations are not attributable to age-related differences among patients.

Together, these findings demonstrate that regional trophic levels, rather than global trophic coherence, are strongly associated with behavioral responsiveness following severe brain injury.

### Dynamic trophic states reveal pathological state trapping

To determine whether trophic hierarchy reorganizes dynamically over time, recurrent whole-brain dynamic trophic states were identified using k-means clustering of instantaneous phase-coupling patterns (Fig. 3). Elbow analysis supported a seven-state solution (Supplementary Fig. 1), from which two biologically distinct states emerged as the principal focus of subsequent analyses: a healthy/integrative state (raw State 3) and a pathological/UWS-enriched state (raw State 7). Relative to the healthy/integrative state, the pathological/UWS-enriched state exhibited higher regional trophic levels across frontal (0.270 versus 0.238), thalamic (0.380 versus 0.298), and insular (0.278 versus 0.163) systems (Fig. 3B). Global trophic coherence differed only modestly between the two states (healthy: 0.00733; pathological: 0.00684), indicating that the principal distinction reflected regional redistribution of trophic hierarchy rather than a marked increase in whole-network trophic coherence (Fig. 3B–D).

**Figure 3.**
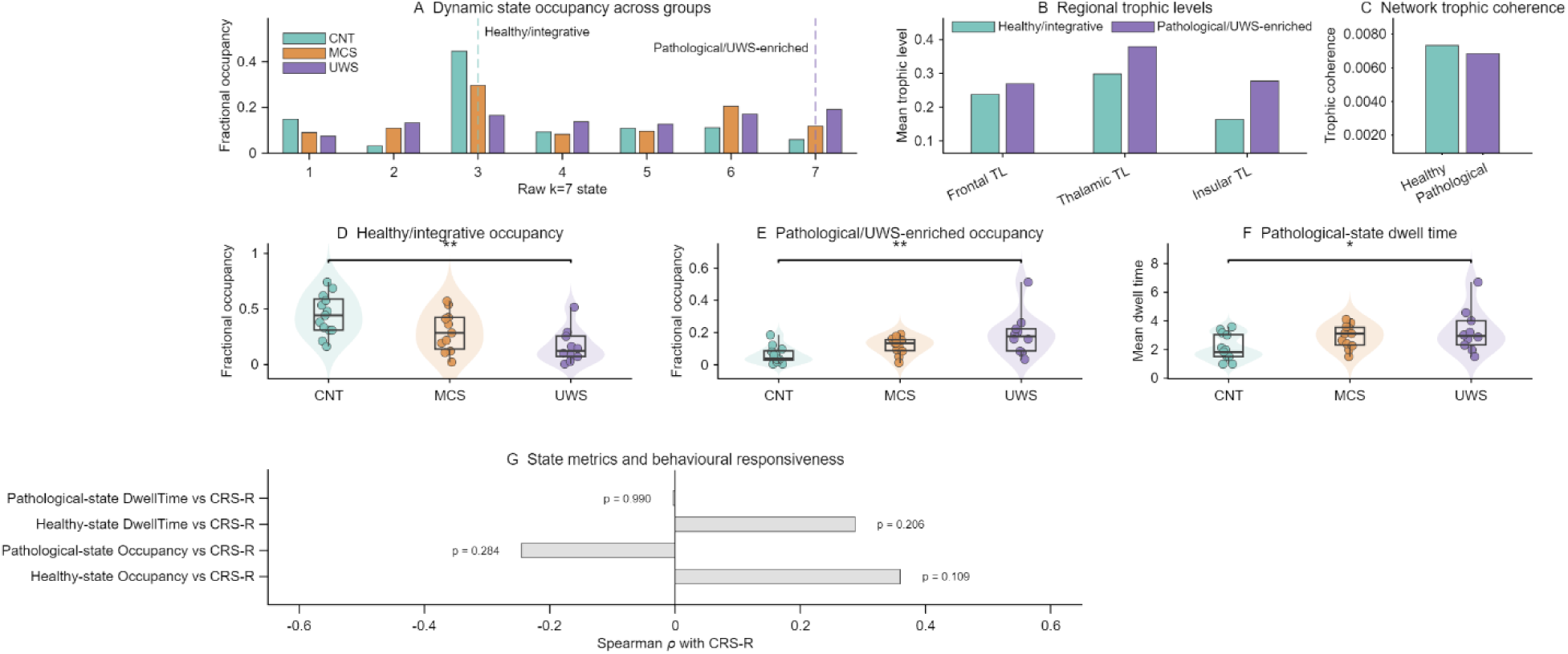
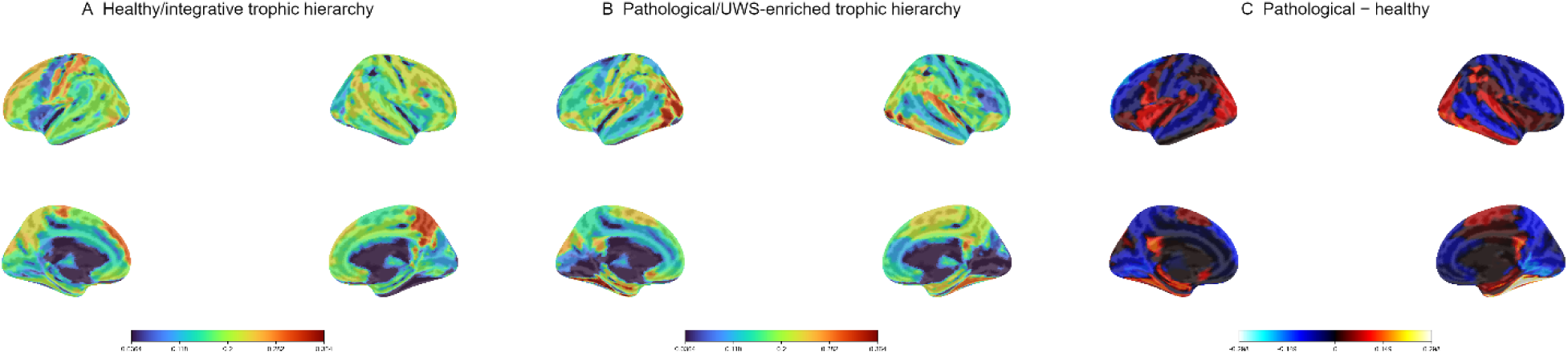
| Dynamic trophic-state organization distinguishes healthy and pathological brain dynamics. **a**, Mean fractional occupancy of all seven dynamic trophic states in CNT, MCS and UWS. State 3 was identified as the healthy/integrative state, whereas State 7 represented the pathological/UWS-enriched state. **b**, Regional trophic organization of the healthy/integrative and pathological/UWS-enriched states. Relative to the healthy state, the pathological state exhibited higher frontal (0.270 versus 0.238), thalamic (0.380 versus 0.298), and insular (0.278 versus 0.163) trophic levels. **c**, Global trophic coherence was higher for the healthy state (0.007329) than for the pathological state (0.006838). **d**, Healthy/integrative-state occupancy differed significantly across diagnostic groups (one-way ANOVA, *F*(2,31)=7.49, *P*=0.00222, η^2^=0.326). **e**, Pathological/UWS-enriched-state occupancy also differed significantly across groups (*F*(2,31)=6.79, *P*=0.00357, η^2^=0.305). **f**, Dwell time within the pathological/UWS-enriched state increased across diagnostic severity (*F*(2,31)=3.66, *P*=0.0373, η^2^=0.191). **g**, None of the state metrics showed significant correlations with CRS-R (healthy occupancy ρ=0.35, *P*=0.109; pathological occupancy ρ=−0.27, *P*=0.284; healthy dwell time ρ=0.29, *P*=0.206; pathological dwell time ρ≈0.00, *P*=0.990). **Figure 3H | Anatomical localization of healthy and pathological dynamic trophic hierarchies**. **a**, Cortical distribution of regional trophic levels for the healthy/integrative state (raw State 3). **b**, Cortical distribution of regional trophic levels for the pathological/UWS-enriched state (raw State 7). **c**, Difference map (pathological minus healthy), highlighting regions with increased trophic level in the pathological state. Surface renderings were generated from state-specific effective connectivity using the same visualization pipeline applied throughout the manuscript.

The temporal expression of these dynamic trophic states differed systematically across diagnostic groups. Fractional occupancy of the healthy/integrative state progressively decreased from healthy controls (0.446 ± 0.181) to minimally conscious state patients (0.297 ± 0.148) and unresponsive wakefulness syndrome patients (0.165 ± 0.086) (one-way ANOVA, *F*(2,31) = 7.500, *P* = 0.002, η^2^ = 0.326). Conversely, occupancy of the pathological/UWS-enriched state increased across the same groups (CNT: 0.060 ± 0.052; MCS: 0.118 ± 0.076; UWS: 0.191 ± 0.120; *F*(2,31) = 6.803, *P* = 0.004, η^2^ = 0.305). These complementary occupancy profiles indicate a progressive redistribution of brain dynamics away from the healthy/integrative state and toward the pathological/UWS-enriched state with increasing impairment of consciousness.

The healthy/integrative state also exhibited significantly longer dwell times in healthy controls than in patient groups (CNT: 8.236 ± 2.637; MCS: 5.716 ± 2.315; UWS: 4.262 ± 2.200; *F* (2,31) = 9.678, *P* < 0.001, η^2^ = 0.385), whereas dwell time within the pathological/UWS-enriched state increased across diagnostic groups (CNT: 2.105 ± 0.892; MCS: 2.925 ± 1.471; UWS: 3.537 ± 1.864; *F*(2,31) = 3.719, *P* = 0.037, η^2^ = 0.191). Together, these findings indicate that disorders of consciousness are characterized by reduced expression of the healthy/integrative state together with increased occupancy and prolonged dwell time within the pathological/UWS-enriched state. These state-specific trophic-level alterations localized predominantly to frontal association cortex together with thalamic and insular regions, as illustrated by the corresponding trophic-level surface renderings (Fig. 5).

Neither healthy-state nor pathological-state occupancy or dwell time correlated significantly with behavioral responsiveness (all *P* > 0.10; Fig. 3H), suggesting that these dynamic measures primarily distinguish diagnostic groups rather than explaining continuous variation in CRS-R scores within the patient cohort.

### Reduced whole-brain dynamical flexibility accompanies pathological trophic-state trapping

To determine whether pathological trophic-state trapping was accompanied by broader alterations in whole-brain dynamics, metastability and global synchrony were quantified using the Kuramoto order parameter (Fig. 4). Metastability progressively decreased across the spectrum of consciousness, from healthy controls (0.208 ± 0.023) to minimally conscious state patients (0.183 ± 0.032) and unresponsive wakefulness syndrome patients (0.164 ± 0.032), demonstrating a significant group effect (one-way ANOVA, *F*(2,31) = 6.862, *P* = 0.003, η^2^ = 0.307). Post hoc analysis revealed significantly lower metastability in UWS than in healthy controls (*P* = 0.003), whereas differences between CNT and MCS (*P* = 0.113) and between MCS and UWS (*P* = 0.261) did not reach statistical significance.

**Figure 4.**
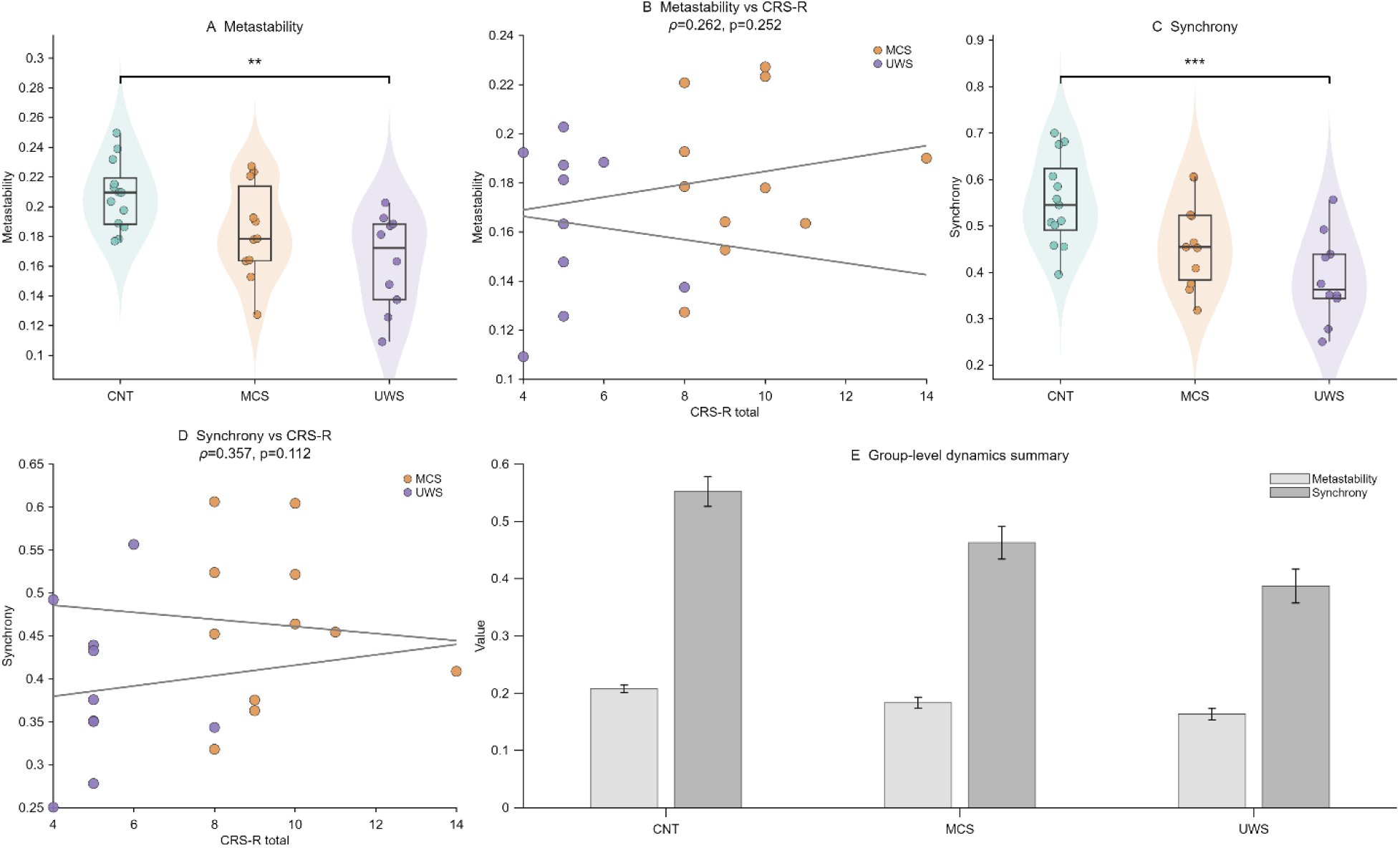
| Reduced metastability and synchrony characterize disorders of consciousness. **a**, Whole-brain metastability differed significantly across groups (one-way ANOVA, *F*(2,31)=6.86, *P*=0.00341, η^2^=0.307). **b**, Metastability was not significantly associated with CRS-R (Spearman ρ=0.262, *P*=0.252). **c**, Whole-brain synchrony differed significantly across groups (*F*(2,31)=8.80, *P*=9.42×10–^4^, η^2^=0.362). **d**, Synchrony was not significantly correlated with CRS-R (ρ=0.357, *P*=0.112). **e**, Group means ± s.e.m. summarizing metastability and synchrony across diagnostic groups.

**Figure 5.**
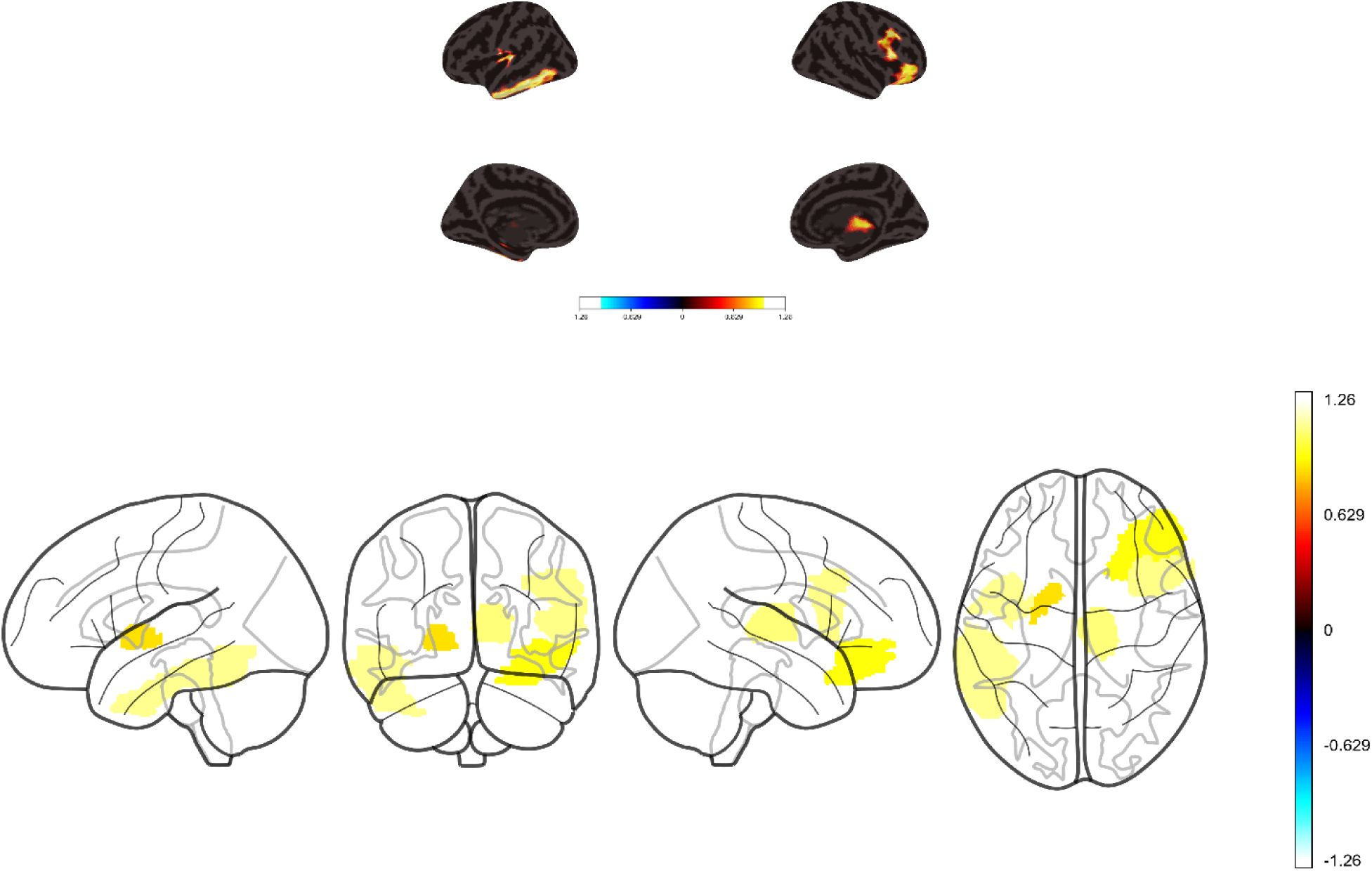
| Exploratory regional trophic-level alterations between healthy controls and unresponsive wakefulness syndrome. Significant uncorrected CNT–UWS differences in regional trophic level projected onto cortical surfaces and complementary glass-brain views. Results are displayed for anatomical interpretation only and were not corrected for multiple comparisons. These analyses are exploratory and identify candidate regions for future confirmatory studies.

Global synchrony exhibited a similar progressive reduction, decreasing from 0.552 ± 0.095 in healthy controls to 0.463 ± 0.095 in MCS and 0.387 ± 0.094 in UWS patients (one-way ANOVA, *F*(2,31) = 8.797, *P* < 0.001, η^2^ = 0.362). Post hoc testing demonstrated significantly reduced synchrony in UWS relative to healthy controls (*P* < 0.001), whereas the remaining pairwise comparisons were not statistically significant (CNT versus MCS: *P* = 0.069; MCS versus UWS: *P* = 0.173).

Neither metastability (Spearman ρ = 0.262, *P* = 0.252) nor global synchrony (ρ = 0.357, *P* = 0.112) correlated significantly with behavioral responsiveness across patients. Together, these findings indicate that disorders of consciousness are associated with reduced whole-brain dynamical flexibility and global coordination, consistent with the increased occupancy and prolonged persistence of the pathological/UWS-enriched trophic state observed in the dynamic trophic-state analysis.

### Robustness across clustering resolutions

To determine whether the identified dynamic trophic states depended on the selected clustering resolution, the complete analysis was repeated across clustering solutions ranging from *k* = 5 to *k* = 9 (Supplementary Figs. 2–4). Although the numerical labels assigned to individual states varied across clustering solutions, the pathological/UWS-enriched state consistently re-emerged. Across all values of *k*, this state consistently exhibited elevated frontal, thalamic, and insular trophic levels together with progressively greater occupancy in patients with unresponsive wakefulness syndrome. The regional hierarchical profile of the pathological/UWS-enriched state therefore remained stable across clustering resolutions despite variation in the numerical state labels.

### Convergent validity under an independent hidden Markov model

To determine whether the pathological/UWS-enriched dynamic trophic state reflects a genuine feature of DOC dynamics rather than an artefact of the LEiDA/k-means pipeline, we repeated the dynamic-state analysis using a Gaussian hidden Markov model (HMM) fitted directly to the regional BOLD time series (Supplementary Table 3, Supplementary Table 4, Supplementary Fig. 5).

The healthy-state finding replicated robustly under this independent framework. At the primary *K* = 7 solution, the state with highest occupancy in healthy controls showed markedly reduced occupancy, dwell time, and self-transition probability in patients (occupancy: *F*(2,31) = 44.403, *P* = 7.94 × 10–^10^, η^2^ = 0.741; dwell time: *F*(1,15) = 5.869, *P* = 0.029; self-transition probability: *F*(1,15) = 5.007, *P* = 0.041; both restricted to CNT versus MCS, since no UWS patient ever occupied this state). Inspection of subject-level occupancy revealed an almost categorical pattern: every healthy control showed substantial occupancy of this state, whereas all ten UWS patients showed zero occupancy, and only 4 of 11 MCS patients ever occupied this state at all.

The pathological-state finding was also well supported, and helped specify which feature of its dynamics is most robust. Occupancy of the state with highest mean occupancy in UWS patients differed significantly across groups at four of five clustering resolutions (*K* = 5, 6, 8, 9; all *P* < 0.03) and showed the same direction of effect at *K* = 7 (*F*(2,31) = 2.520, *P* = 0.097, η^2^ = 0.140), the resolution matching the manuscript’s primary LEiDA solution, indicating that increased occupancy of this configuration is a reasonably consistent feature of DOC dynamics across model choices. Dwell time of this state reached significance only at *K* = 5 (*P* = 0.036) and was non-significant at all other resolutions (*K* = 6–9, *P* > 0.18). The self-transition probability of the pathological state did not differ significantly across groups at any clustering resolution tested (*K* = 5–9, all *P* > 0.15; at the primary K=7 solution, both the dwell-time and self-transition comparisons for this state reduce to a comparison between MCS and UWS patients only, since no control participant ever occupied it — a further, independent illustration of the same access-based pattern). Taken together, these results indicate that entry into the pathological configuration, rather than an increase in its intrinsic stability once entered, is the more robust and reproducible signature of altered dynamics in DOC.

Because occupancy of both states was strongly bimodal across subjects, we tested whether the probability of ever occupying a given state, rather than its continuous occupancy, differentiated diagnostic groups. This categorical measure showed a clear and statistically robust gradient: no healthy control ever occupied the pathological-candidate state (0/13), compared with 3 of 11 MCS patients (27%) and 7 of 10 UWS patients (70%) (χ^2^(2) = 13.376, *P* = 0.0012; monotonic trend across diagnostic severity, Spearman ρ = 0.619, *P* = 1.0 × 10–^4^). The converse pattern held for the healthy-candidate state (100% of CNT, 36% of MCS, and 0% of UWS patients ever occupied this state).

Pathological-state occupancy under the HMM showed a similar direction of association with CRS-R as under LEiDA, though again non-significant (ρ = −0.406, *P* = 0.068, *N* = 21). Correlations involving dwell time and self-transition probability of either state were not reliably interpretable, as a majority of patients never occupied one or both states at all, restricting these analyses to small subsets of the patient sample (*N* = 4–10).

## Discussion

The present study investigated the relationship between hierarchical brain organization, large-scale brain dynamics, and behavioral responsiveness in disorders of consciousness. By combining trophic-level analysis, trophic coherence, dynamic state identification, metastability, and behavioral assessment, we identified a previously unrecognized dissociation between static and dynamic hierarchy. Whereas higher static fronto-thalamic trophic levels were associated with better behavioral responsiveness, as measured by the Coma Recovery Scale–Revised (CRS-R), the pathological hyper-hierarchical state was associated with more severe impairment of consciousness. Patients with severe disorders of consciousness exhibited increased occupancy and prolonged residence within a pathological/UWS-enriched state (hereafter, pathological hyper-hierarchical state) characterized by elevated frontal, thalamic, and insular trophic levels, reduced metastability, and altered global synchrony. Together, these findings suggest that conscious awareness depends not only on hierarchical organization itself, but also on the capacity to flexibly transition between distinct hierarchical brain states.

Our findings extend contemporary theories of consciousness by demonstrating that hierarchical organization plays fundamentally different roles across temporal scales. At the static level, higher frontal and thalamic trophic levels were associated with better behavioral responsiveness, and together with insular trophic levels explained approximately two-thirds of the variance in CRS-R scores. These findings indicate that the regional organization of directed brain networks is strongly related to residual conscious function and are consistent with extensive evidence implicating fronto-thalamic systems in conscious access, arousal regulation, and recovery following severe brain injury^1–7^.

The thalamus occupies a central position in mechanistic accounts of consciousness. Schiff and colleagues proposed that recovery of consciousness depends on the restoration of large-scale thalamocortical interactions^48,51^, while neuroimaging studies have consistently identified thalamic dysfunction as a hallmark of disorders of consciousness^17–21^. Medial frontal regions likewise constitute key components of the frontoparietal networks implicated in global broadcasting, executive integration, and conscious report^7,10,11,56^. Within this framework, the positive association between fronto-thalamic trophic hierarchy and CRS-R scores suggests that maintenance of directed hierarchical organization within these systems supports residual conscious function following severe brain injury.

In contrast to the static analyses, the dynamic results indicate that hierarchical organization becomes maladaptive when concentrated within recurrent brain states. Rather than exhibiting a generalized loss of hierarchical structure, patients with severe disorders of consciousness showed increased occupancy and prolonged residence within a pathological hyper-hierarchical state characterized by elevated frontal, thalamic, and insular trophic levels together with increased trophic coherence. This dissociation between static and dynamic hierarchy constitutes one of the principal findings of the present study. If hierarchical organization were uniformly beneficial, one would expect both static and dynamic measures to show similar relationships with conscious function. Instead, our findings suggest that the functional significance of hierarchy depends critically on its temporal organization.

At the static level, higher regional trophic hierarchy may support efficient large-scale communication and the fronto-thalamic interactions necessary for conscious processing. At the dynamic level, however, excessive concentration of hierarchical organization within recurrent brain states may restrict transitions between alternative network configurations. Conscious awareness may therefore depend not simply on the presence of hierarchical organization, but on the capacity to flexibly redistribute hierarchical control over time.

Static hierarchy may provide an organizational scaffold that supports efficient large-scale communication. In contrast, excessive concentration of hierarchical organization within recurrent dynamic states may limit exploration of alternative network configurations. From this perspective, the pathological dynamics observed in disorders of consciousness resemble excessive attractor stability rather than network disintegration^13–16,35–41^. This interpretation is supported by the identification of a pathological hyper-hierarchical state characterized by elevated trophic coherence together with increased frontal, thalamic, and insular trophic levels. Rather than reflecting a loss of hierarchical organization, this state represents an unusually ordered configuration in which directed interactions become concentrated within a restricted dynamical regime. Its progressively greater occupancy and prolonged dwell time in patients with more severe disorders of consciousness suggest that the principal abnormality is not the disappearance of hierarchical structure, but pathological trapping within a hyper-hierarchical brain state.

This interpretation is consistent with contemporary dynamical systems accounts of whole-brain function. Computational models developed by Deco, Kringelbach, Cabral, and colleagues propose that healthy brain activity occupies a metastable regime characterized by continual transitions among partially synchronized network configurations^18–22^. Within this framework, consciousness emerges from a dynamic balance between integration and segregation that enables flexible exploration of large-scale brain states^23–26^. Our findings extend this perspective by suggesting that disorders of consciousness are characterized not by a reduction in the number of available brain states, but by pathological trapping within an excessively stable hyper-hierarchical configuration. The progressive increase in occupancy and dwell time of the pathological hyper-hierarchical state, together with reduced metastability, is consistent with a shift toward a less flexible dynamical regime in which transitions between alternative network configurations become increasingly constrained.

The whole-brain dynamical analyses provide independent support for this interpretation. Consistent with previous studies of disorders of consciousness, both metastability and global synchrony were significantly reduced in patients with more severe impairment of consciousness. Metastability reflects the temporal variability of whole-brain synchronization and is widely interpreted as an index of dynamical flexibility^18,21,27^. The observed reduction in metastability therefore supports the view that pathological hyper-hierarchical state trapping is accompanied by a diminished capacity to transition between alternative large-scale network configurations. Likewise, reduced global synchrony is consistent with previous work suggesting that conscious processing depends not on maximal synchronization or desynchronization, but on an intermediate dynamical regime that supports both large-scale integration and functional differentiation^28–31^. Together, these findings indicate that pathological hyper-hierarchical organization emerges within a broader context of impaired whole-brain dynamical flexibility.

The pathological hyper-hierarchical state was characterized by elevated trophic levels across a distributed fronto-thalamo-limbic network, including the thalamus, basal ganglia, insula, amygdala, parahippocampal cortex, supplementary motor area, and medial frontal cortex. These regions largely overlap with systems implicated in arousal regulation, salience processing, motivational control, and large-scale network coordination^53–55^. Rather than suggesting diffuse disruption of brain organization, this anatomical distribution is consistent with the emergence of a coherent hyper-hierarchical network that may act as a dynamical bottleneck, restricting transitions to more flexible large-scale brain states.

The prominent involvement of the insula is particularly noteworthy. Whereas lower static insular trophic levels were associated with better behavioral responsiveness, the pathological hyper-hierarchical state exhibited markedly elevated insular hierarchy, suggesting that the functional role of this region differs across temporal scales. As a central hub of the salience network, the insula has been proposed to mediate switching between internally and externally directed modes of brain activity^54,55^. Excessive hierarchical dominance within this system may therefore contribute to pathological state stabilization by repeatedly biasing large-scale brain dynamics toward a restricted hyper-hierarchical configuration.

The present findings also have implications for broader theories of hierarchical brain organization. Recent work using fluctuation–dissipation approaches has similarly reported disrupted hierarchical organization in disorders of consciousness, despite employing a fundamentally different measure of hierarchy.^49,50^. Predictive processing accounts propose that cortical function is fundamentally hierarchical, with higher levels generating predictions and lower levels transmitting prediction errors^38,39,59^. Within these frameworks, hierarchy is not inherently beneficial or detrimental; rather, adaptive brain function depends on flexible interactions across hierarchical levels. Although the present analyses were not designed to directly test predictive-processing models, the observation that hierarchical organization becomes concentrated within pathological hyper-hierarchical states is broadly consistent with the idea that impaired consciousness reflects a loss of flexibility in hierarchical information processing rather than disruption of hierarchy itself.

Throughout this manuscript we have used the language of “trapping” to describe the joint pattern of increased occupancy and dwell time observed under LEiDA; the analysis below indicates that the entry/occupancy component of this pattern is the more robust and independently reproducible one. To assess whether the pathological/UWS-enriched state reflects a genuine feature of DOC dynamics rather than an artefact of the LEiDA/k-means clustering pipeline, we repeated the dynamic-state analysis using an independent framework, a Gaussian hidden Markov model fitted directly to the regional BOLD time series^26,69^ (Supplementary Table 3, Supplementary Table 4, Supplementary Fig. 5). This convergent-validity analysis provided partial support for our central claims. The reduction in occupancy of a healthy/integrative configuration in patients replicated robustly, and subject-level inspection revealed an even more categorical pattern than under LEiDA: UWS patients almost never occupied this configuration at all, rather than occupying it with reduced frequency. Occupancy of a UWS-enriched pathological configuration also increased with diagnostic severity at most clustering resolutions, and the probability of ever entering this configuration, rather than the amount of time spent within it once entered, showed a particularly robust gradient across diagnostic groups. The self-transition probability of this state, however, did not differ across groups at any clustering resolution tested. Read together with the occupancy and presence/absence results, this convergent-validity analysis refines rather than undermines the manuscript’s central claim: across two independent dynamic-state frameworks, patients with more severe impairment are consistently more likely to ever enter an aberrant, UWS-enriched configuration, and this increased accessibility — rather than a change in the configuration’s intrinsic stability once entered — is the most robust and reproducible signature of altered dynamics in DOC. This distinction may be neurobiologically meaningful rather than purely statistical: a configuration that becomes progressively easier to enter, without a corresponding increase in the difficulty of leaving it, is more consistent with a widened basin of attraction — one reachable from a broader range of preceding brain states — than with a deepened or more stable attractor. A more mundane contribution cannot be excluded, however: most patients who entered the pathological state at all did so only briefly (Supplementary Table 3), so self-transition probability was estimated from relatively few within-state transitions for many subjects, which may have limited our power to detect a true persistence difference had one been present. We therefore characterize the pathological dynamic state as one that becomes increasingly accessible, and increasingly likely to be entered at all, as impairment of consciousness becomes more severe, while treating its characterization as unusually persistent or self-reinforcing once entered as a secondary feature that is less strongly supported by the present data.

More broadly, our findings challenge the traditional view of disorders of consciousness as simple disconnection syndromes. Rather than observing a global collapse of hierarchical organization, we found that regional trophic hierarchy remained behaviorally relevant while pathological hyper-hierarchical configurations increasingly dominated large-scale brain dynamics. This distinction may help reconcile apparently conflicting observations in the consciousness literature, where some studies report reduced functional integration whereas others describe preserved— or even elevated—coordination within specific large-scale networks^22–25^. Our results suggest that these findings are not necessarily contradictory: disorders of consciousness may simultaneously exhibit preserved regional hierarchical organization and pathological concentration of that hierarchy within restricted dynamical states.

Several limitations should be acknowledged. First, the study relied on resting-state fMRI, which provides an indirect measure of neural activity and has limited temporal resolution. Second, trophic hierarchy represents one of several approaches for quantifying directed network organization. Recent work using complementary hierarchy measures suggests that hierarchical abnormalities may generalize across analytical frameworks, but future studies should determine whether the dissociation between static and dynamic hierarchy identified here extends across alternative hierarchy metrics.^49,50^ Third, although the pathological hyper-hierarchical state emerged consistently across clustering resolutions, dynamic state identification inevitably depends on methodological choices, including temporal resolution and clustering strategy. Finally, the cross-sectional design precludes causal inference regarding the role of trophic hierarchy in recovery and longitudinal changes in consciousness. Fifth, although the association between frontal and insular trophic levels and CRS-R survived adjustment for age, leave-one-out re-estimation indicated that the frontal association was sensitive to individual patients, in some iterations falling below the significance threshold, whereas the insular association remained stable throughout (Supplementary Table 2, Supplementary Fig. 4); this association should therefore be interpreted with some caution pending replication in larger cohorts. Sixth, an independent hidden Markov model analysis confirmed that patients with more severe impairment are consistently more likely to ever enter a pathological dynamic state (Supplementary Table 3, Supplementary Table 4, Supplementary Fig. 5), but did not show a corresponding increase in that state’s self-transition probability. This indicates that increased accessibility of the pathological configuration, rather than an increase in its intrinsic stability once entered, is the better-supported feature of the present findings.

An additional methodological consideration concerns the use of the structural connectivity matrix. As in previous whole-brain modelling studies of disorders of consciousness, the models were initialized using a structural connectivity matrix derived from healthy participants. Given the marked heterogeneity of structural lesions across patients, a normative anatomical scaffold provides a consistent starting point for model optimization. Importantly, this structural connectivity matrix served only as the anatomical prior for the modelling procedure. Participant-specific effective connectivity was subsequently estimated by iteratively optimizing directed interactions to reproduce each individual’s empirical functional connectivity and time-lagged covariance structure, allowing individualized effective connectivity networks to emerge despite the common structural scaffold.

The present findings suggest that disorders of consciousness are characterized not by the collapse of hierarchical organization, but by its pathological concentration within a restricted set of recurrent brain states. Whereas higher static fronto-thalamic trophic hierarchy was associated with better behavioral responsiveness, severe disorders of consciousness were characterized by increased occupancy and prolonged residence within a pathological hyper-hierarchical state exhibiting elevated frontal, thalamic, and insular trophic levels together with reduced metastability and altered global synchrony. These findings suggest that conscious awareness depends not simply on the presence of hierarchical brain organization, but on the capacity to flexibly transition between distinct hierarchical configurations. Pathological trapping within hyper-hierarchical brain states may therefore represent a previously unrecognized dynamical signature of impaired consciousness and a potential target for future mechanistic and therapeutic investigations.

## Notes

### Competing Interest Statement

The authors have declared no competing interest.

