## Supplementary figures for "Hyper-Hierarchical Brain States Are Associated with Disorders of Consciousness"

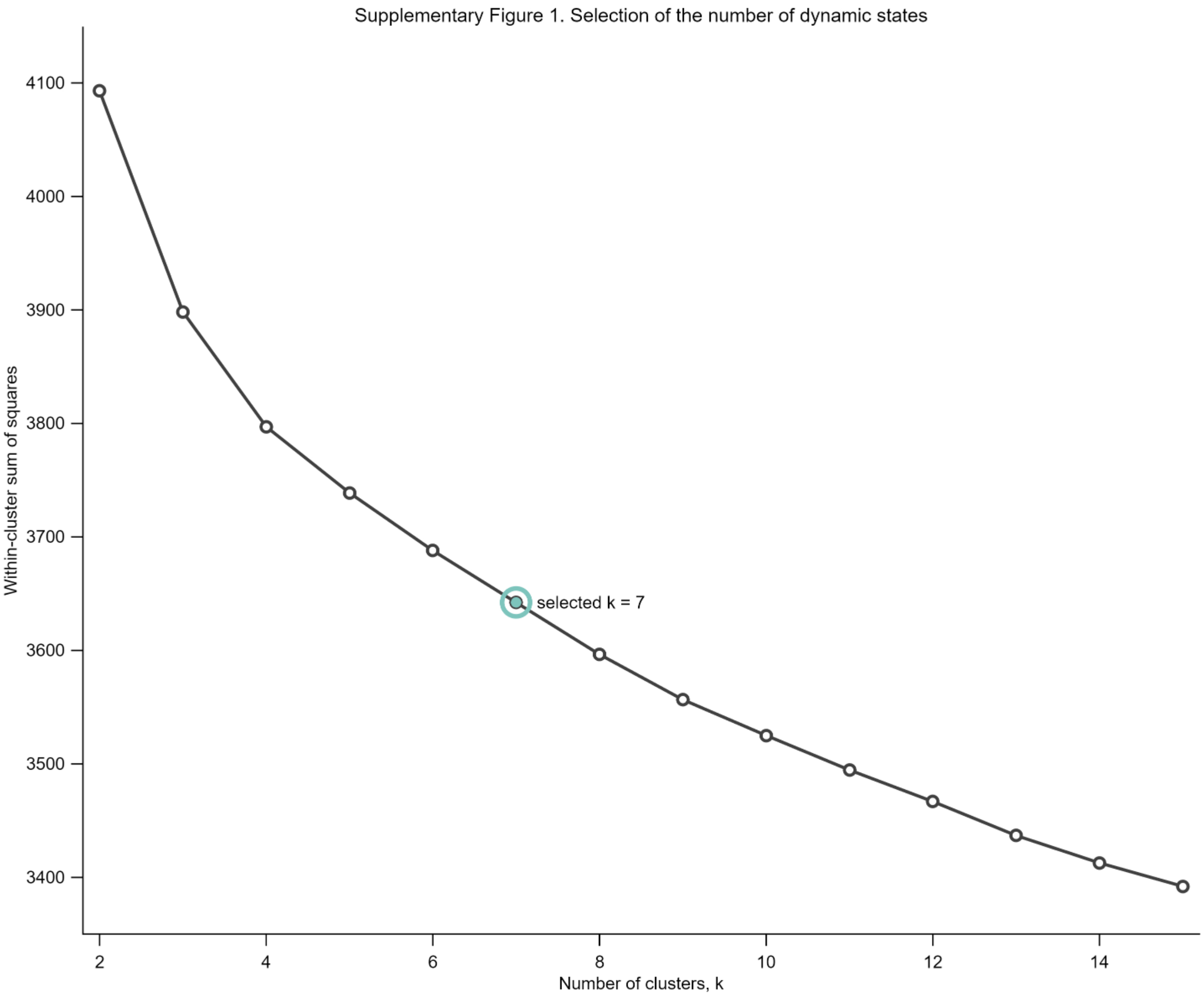


**Supplementary Figure 1 | Selection of the number of dynamic states.**

Elbow analysis showing within-cluster sum of squares for k=2–15 LEiDA clusters. The reduction in clustering error diminished beyond k≈7, supporting the selection of seven dynamic states for all subsequent analyses. The selected solution (k=7) is highlighted.

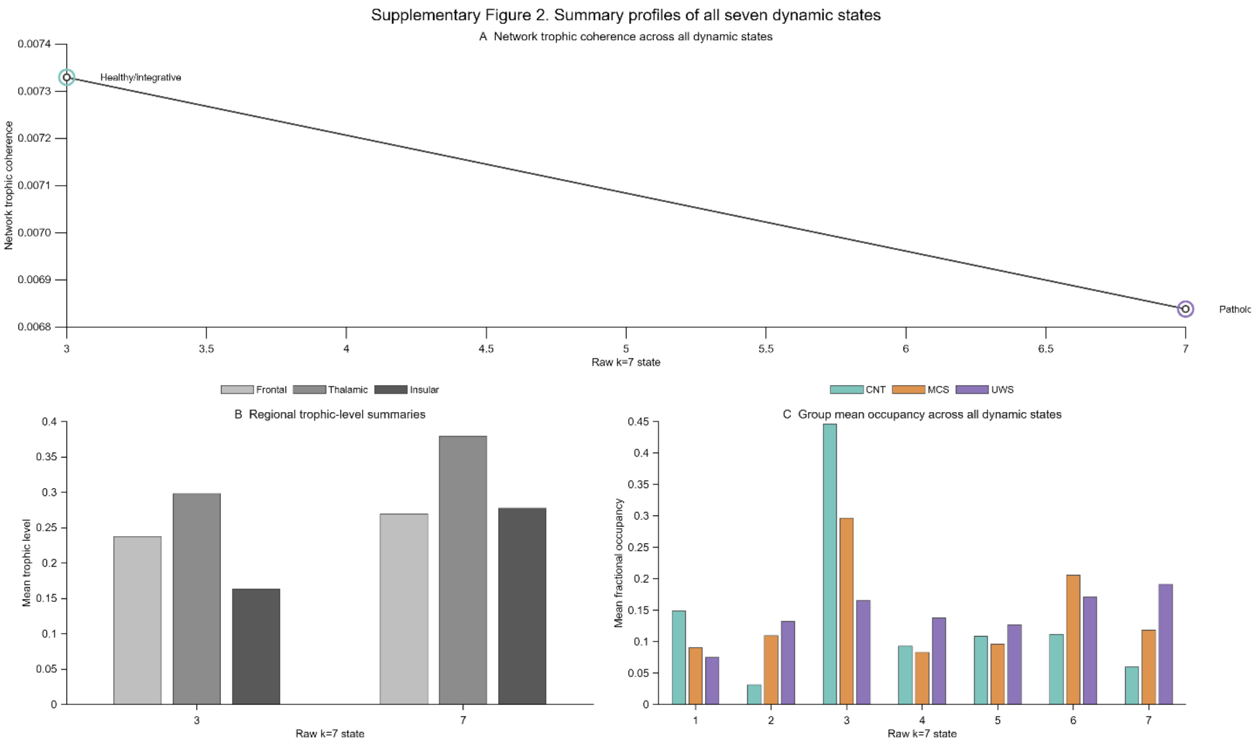


**Supplementary Figure 2 - Summary characteristics of all seven dynamic trophic states.**

States are ordered according to the original LEiDA k = 7 solution. State 3 corresponds to the healthy/integrative state and State 7 to the pathological/UWS-enriched state; States 1, 2, 4, 5 and 6 are shown for completeness but were not the focus of the primary analyses.

**a**, Network trophic coherence for each state.

**b**, Mean frontal, thalamic and insular trophic levels for the healthy/integrative (State 3) and pathological/UWS-enriched (State 7) states.

**c**, Mean fractional occupancy of all seven states in CNT, MCS and UWS, illustrating that the healthy state is preferentially expressed in controls whereas the pathological state is progressively enriched in disorders of consciousness.

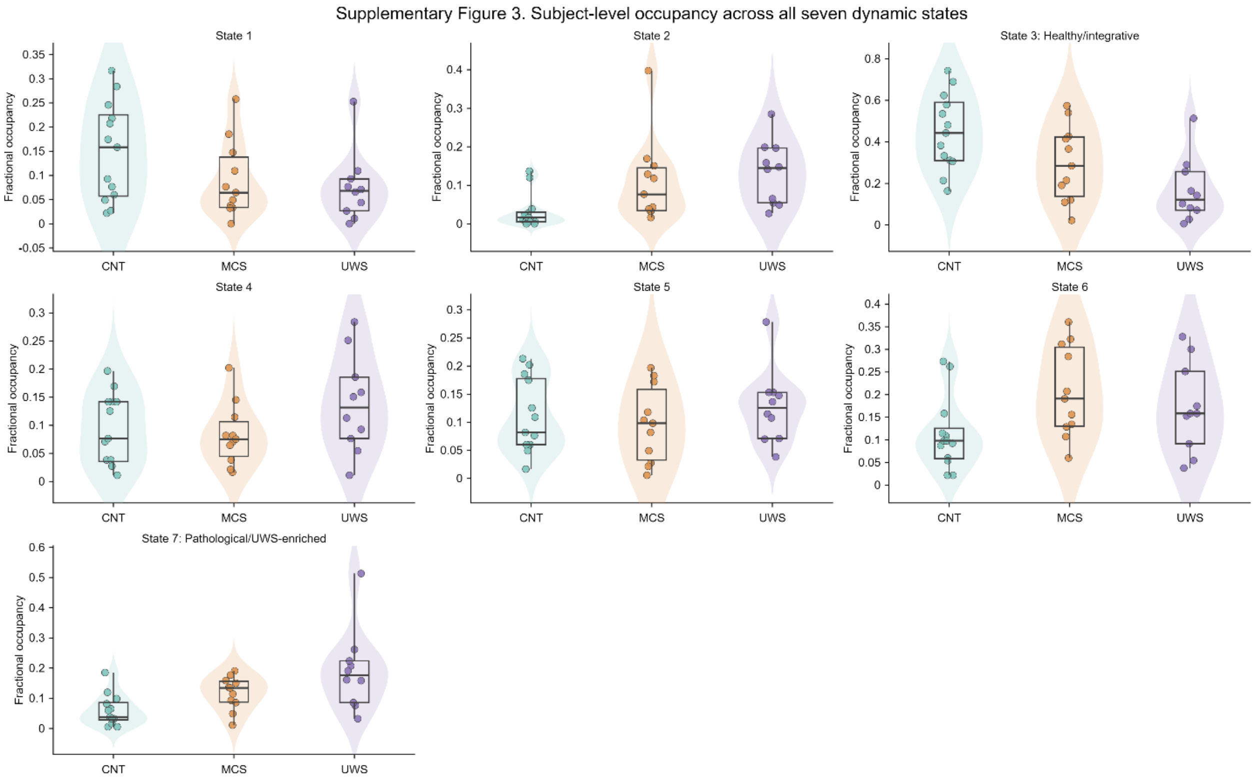


**Supplementary Figure 3 | Subject-level occupancy distributions for all seven dynamic trophic states.**

Violin plots display individual fractional occupancies for each dynamic state across CNT, MCS and UWS participants. Boxplots denote median and interquartile range; points represent individual participants. State 3 corresponds to the healthy/integrative state and State 7 to the pathological/UWS-enriched state.

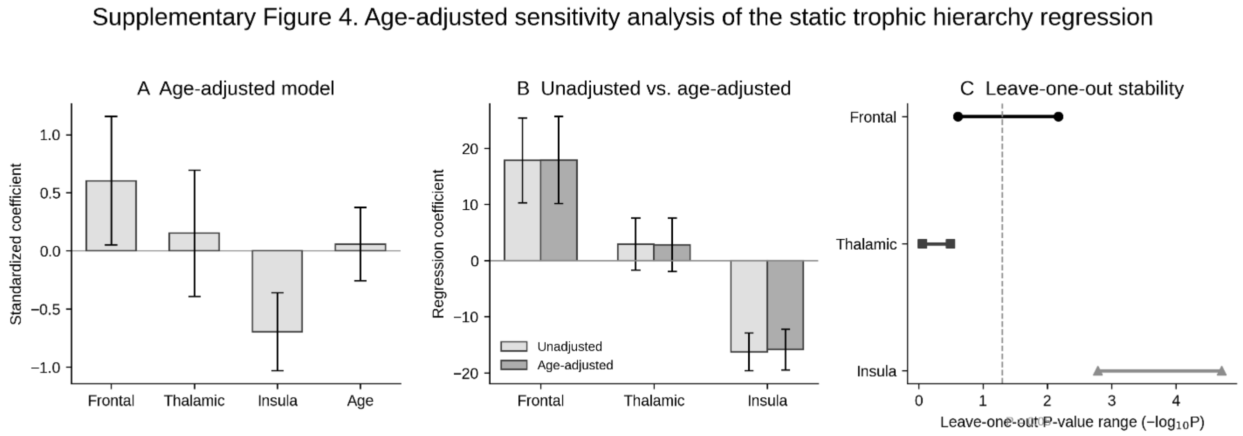


**Supplementary Figure 4 | Age-adjusted sensitivity analysis of the static trophic-hierarchy regression.**

The multiple regression relating regional trophic levels to CRS-R (Fig. 2e) was repeated with participant age included as a covariate (N=21 patients). **a**, Standardized coefficients (± 95% CI) from the age-adjusted model. Age was not associated with CRS-R (β=0.059, P=0.697) and did not improve model fit relative to the unadjusted model ($\Delta R^2$=0.003; F-change(1,16)=0.157, P=0.697). **b**, Unstandardized regression coefficients (± SE) for frontal, thalamic, and insular trophic levels in the unadjusted versus age-adjusted models, illustrating minimal change after age adjustment. **c**, Leave-one-out stability of each predictor's association with CRS-R, showing the range of P-values ($-\log_{10}P$) obtained across 21 leave-one-out iterations of the age-adjusted model. The insular association remained significant across all iterations; the frontal association was more sensitive to individual observations, in some iterations falling below the significance threshold (dashed line).

Supplementary Figure 5. Convergent-validity analysis of the pathological dynamic state using an independent hidden Markov model

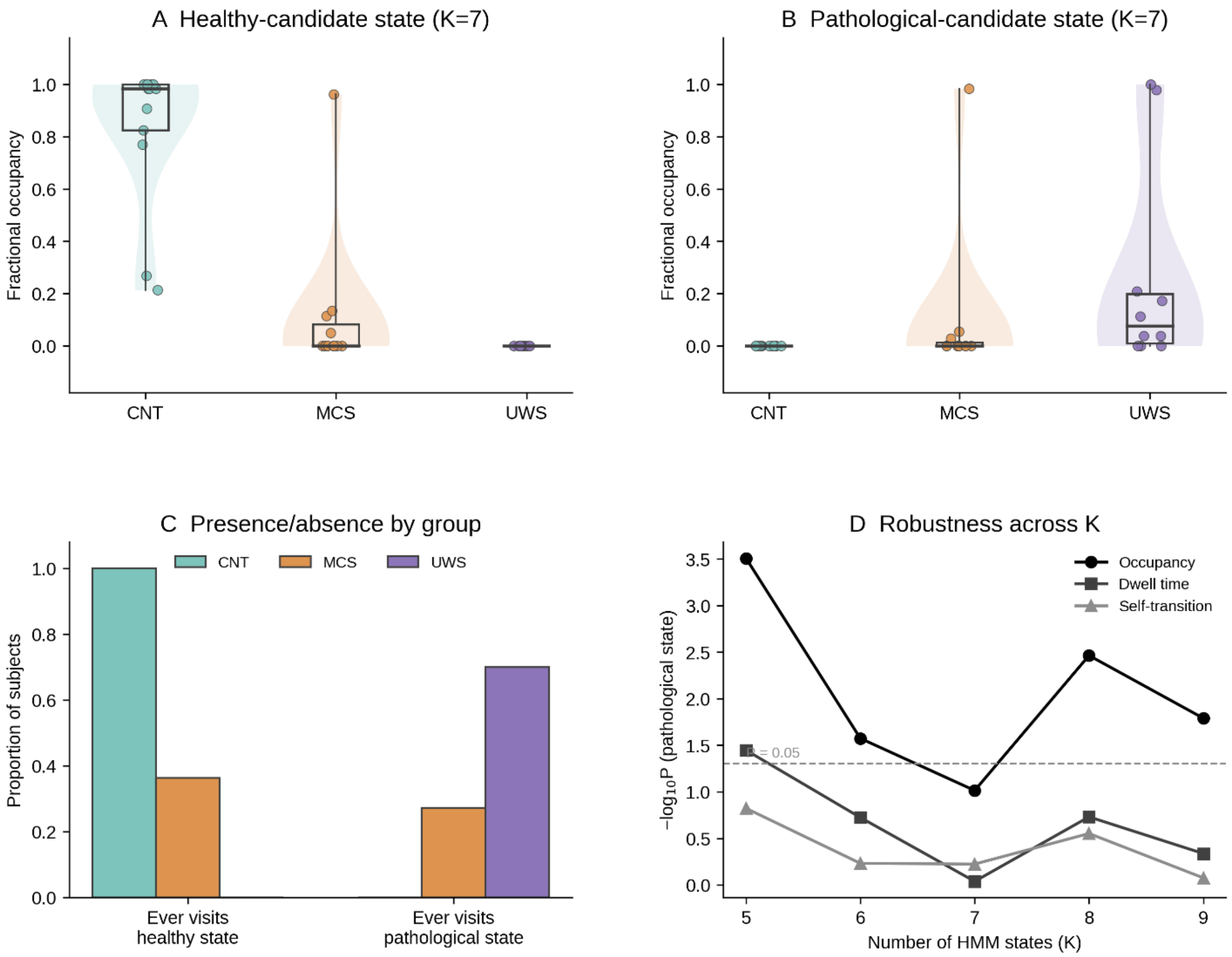

**Supplementary Figure 5 | Convergent-validity analysis of the pathological dynamic state using an independent hidden Markov model.**

A Gaussian hidden Markov model (HMM-MAR) was fitted directly to the band-pass filtered, z-scored regional BOLD time series, independently of the LEiDA/k-means pipeline. **a**, Mean fractional occupancy of the healthy- and pathological-candidate states by diagnostic group at the primary K=7 solution. **b**, Proportion of subjects in each group who ever occupied the healthy- or pathological-candidate state at all (occupancy > 0), reflecting the strongly bimodal distribution of subject-level occupancy. **c**, Robustness of group differences in occupancy, dwell time, and self-transition probability of the pathological-candidate state across clustering resolutions (K=5–9); see Supplementary Table 4 for the corresponding statistics and a note on group sizes. Occupancy differed significantly across groups at four of five resolutions, indicating that entry into this state is a reasonably robust feature of DOC dynamics; dwell time and self-transition probability were less consistent across resolutions and, at K=7, reflect a comparison restricted to MCS and UWS patients only, since no control participant ever occupied this state.

| Raw state | Manuscript label | Description |
|---|---|---|
| 1 | Intermediate | Intermediate dynamic trophic state retained in the final k = 7 solution; no primary group-enriched manuscript interpretation was assigned. |
| 2 | Intermediate | Intermediate dynamic trophic state retained in the final k = 7 solution; no primary group-enriched manuscript interpretation was assigned. |
| 3 | **Healthy/integrative** | Healthy-control-enriched dynamic state with the highest occupancy in controls and an integrative trophic profile; used as the reference healthy dynamic trophic hierarchy. |
| 4 | Intermediate | Intermediate dynamic trophic state retained in the final k = 7 solution; no primary group-enriched manuscript interpretation was assigned. |
| 5 | Intermediate | Intermediate dynamic trophic state retained in the final k = 7 solution; no primary group-enriched manuscript interpretation was assigned. |
| 6 | Intermediate | Intermediate dynamic trophic state retained in the final k = 7 solution; no primary group-enriched manuscript interpretation was assigned. |
| 7 | **Pathological/UWS-enriched** | UWS-enriched dynamic state with a pathological trophic profile and increased regional trophic levels; used as the pathological dynamic trophic hierarchy. |

**Supplementary Table 1 | Mapping between raw LEiDA cluster indices and manuscript terminology.**

Correspondence between the original k=7 LEiDA state numbering and the terminology adopted throughout the manuscript. States 3 and 7 were identified as the healthy/integrative and pathological/UWS-enriched dynamic trophic hierarchies, respectively, whereas the remaining states are retained using their raw numbering because they were not the focus of the primary analyses.

**Supplementary Table 2 | Age-adjusted static trophic-hierarchy regression.**

Coefficients from the multiple regression relating regional trophic levels to CRS-R (N=21 patients), before and after adjusting for participant age. Unadjusted model: $R^2$=0.677, adjusted $R^2$=0.620, $F(3,17)$=11.867, $P=1.96\times10^{-4}$. Age-adjusted model: $R^2$=0.680, adjusted $R^2$=0.600, $F(4,16)$=8.499, $P=7.09\times10^{-4}$. Adding age did not significantly improve model fit (F-change(1,16)=0.157, P=0.697). Frontal and insular trophic levels remained independent predictors of CRS-R after age adjustment.

| Predictor | Unadjusted β (SE) | Unadjusted P | Age-adjusted β (SE) | Age-adjusted P |
|---|---|---|---|---|
| Frontal | 17.821 (7.541) | 0.0303 | 17.897 (7.737) | 0.0344 |
| Thalamic | 2.910 (4.603) | 0.5356 | 2.783 (4.732) | 0.5646 |
| Insula | −16.284 (3.363) | $1.53\times10^{-4}$ | −15.865 (3.608) | $4.50\times10^{-4}$ |
| Age | — | — | 0.010 (0.025) | 0.6971 |

**Supplementary Table 3 | Hidden Markov model state statistics at K=7.**

One-way ANOVA and Spearman correlation with CRS-R (patients only) for the healthy- and pathological-candidate HMM states at the primary K=7 solution. Occupancy is defined for all subjects and compared across all three groups (df=2,31). Dwell time and self-transition probability are undefined for subjects who never occupy a state at all; because no UWS patient ever occupied the healthy state and no CNT participant ever occupied the pathological state, these comparisons are effectively two-group tests (CNT vs. MCS, df=1,15, for the healthy state; MCS vs. UWS, df=1,8, for the pathological state) rather than three-group ANOVAs, as indicated in the Metric column. N for CRS-R correlations reflects the number of patients with non-zero occupancy of the relevant state.

| Metric | Group F | Group P | $\eta^2$ | CRS-R ρ | CRS-R P | N |
|---|---|---|---|---|---|---|
| Healthy occupancy | 44.403 | $7.94\times10^{-10}$ | 0.741 | 0.325 | 0.151 | 21 |
| Pathological occupancy | 2.520 | 0.0968 | 0.140 | −0.406 | 0.068 | 21 |
| Healthy dwell time (CNT vs MCS) | 5.869 | 0.0285 | 0.281 | 0.000 | 1.000 | 4 |
| Pathological dwell time (MCS vs UWS) | 0.012 | 0.9158 | 0.001 | 0.052 | 0.886 | 10 |
| Healthy self-transition P (CNT vs MCS) | 5.007 | 0.0409 | 0.250 | 0.000 | 1.000 | 4 |
| Pathological self-transition P (MCS vs UWS) | 0.302 | 0.5975 | 0.036 | 0.425 | 0.221 | 10 |

**Supplementary Table 4 | Robustness of pathological-state HMM group differences across K=5–9.**

One-way ANOVA for occupancy, dwell time, and self-transition probability of the pathological-candidate state, repeated across HMM clustering resolutions K=5–9. Occupancy is defined for all subjects and compared across all three groups (df=2,31) at every K. As at K=7 (Supplementary Table 3), no control participant occupied the pathological-candidate state at K=7, and dwell time and self-transition probability are undefined for subjects who never occupy a state; the same collapse to an effectively two-group (MCS vs. UWS) comparison likely applies at other resolutions, though group-wise sample sizes were not separately verified for K=5, 6, 8, and 9. Occupancy differed significantly across groups at four of five resolutions (K=5,6,8,9), supporting increased entry into this state with diagnostic severity; dwell time and self-transition probability were less consistent across resolutions.

| K | Occ. F | Occ. P | Dwell F | Dwell P | SelfTrans F | SelfTrans P |
|---|---|---|---|---|---|---|
| 5 | 10.602 | $3.10\times10^{-4}$ | 5.591 | 0.0358 | 2.361 | 0.1504 |
| 6 | 4.082 | 0.0267 | 1.989 | 0.1888 | 0.316 | 0.5865 |
| 7 | 2.520 | 0.0968 | 0.012 | 0.9158 | 0.302 | 0.5975 |
| 8 | 6.851 | 0.0034 | 1.992 | 0.1858 | 1.289 | 0.2804 |
| 9 | 4.722 | 0.0162 | 0.833 | 0.4604 | 0.177 | 0.8404 |
